# Flexible foraging behaviour increases predator vulnerability to climate change

**DOI:** 10.1101/2021.05.05.442768

**Authors:** Benoit Gauzens, Benjamin Rosenbaum, Gregor Kalinkat, Thomas Boy, Malte Jochum, Susanne Kortsch, Eoin J. O’Gorman, Ulrich Brose

**Affiliations:** EcoNetLab, German Centre for Integrative Biodiversity Research (iDiv) Halle-Jena-Leipzig, Leipzig, Germany; Institute of Biodiversity, Friedrich Schiller University Jena, Jena, Germany; Department of Community and Ecosystem Ecology, Leibniz Institute of Freshwater Ecology and Inland Fisheries (IGB), Berlin, Germany; Experimental Interaction Ecology, German Centre for Integrative Biodiversity Research (iDiv) Halle-Jena-Leipzig, Leipzig, Germany; Leipzig University, Institute of Biology, Leipzig, Germany; Department of Global Change Ecology, Biocenter, University of Würzburg, Würzburg, Germany; Tvärminne Zoological Station, University of Helsinki, Hanko, Finland; School of Life Sciences, University of Essex, Wivenhoe Park, Colchester CO4 3SQ, UK

## Abstract

Higher temperatures are expected to reduce species coexistence by increasing energetic demands. However, flexible foraging behaviour could balance this effect by allowing predators to target specific prey species to maximize their energy intake, according to principles of optimal foraging theory. We test these assumptions using a large dataset comprising 2,487 stomach contents from six fish species with different feeding strategies, sampled across environments with varying prey availability over 12 years in Kiel Bay (Baltic Sea). Our results show that foraging shifts from trait-to density-dependent prey selectivity in warmer and more productive environments. This behavioural change leads to lower consumption efficiency at higher temperature as fish select more abundant but less energetically rewarding prey, thereby undermining species persistence and biodiversity. By integrating this behaviour into dynamic food web models, our study reveals that flexible foraging leads to lower species coexistence and biodiversity in communities under global warming.

## Main text

Ecosystems are experiencing abrupt changes in climatic conditions, making it ever more important to predict and understand how they will respond in the future. Global warming will affect various levels of biological organization; from physiological processes occurring at the individual level^1^ to patterns at macroecological scales^2^. Warming impacts will cascade through these different organizational levels, changing species composition^3^, as well as community and food web structure^4^. By scaling up temperature effects from species physiology to food webs^5^, trophic interactions play a key role in an ecosystem’s response to global warming^6^. Therefore, food web models have become important tools for predicting the future of communities amidst rising temperatures, and understanding the underlying mechanisms^7^. However, current food web models are mostly based on networks composed of static feeding interactions, thereby disregarding flexibility in consumer behaviour in response to varying environmental conditions.

Food web models building on biological processes observed at the level of individual organisms can be used to test mechanisms and generate predictions at the ecosystem level and therefore offer an integrative solution to study temperature effects on communities. For example, Allometric Trophic Networks^8^ (ATNs) quantify effects of body mass and temperature on the biological rates of consumers and resources to predict species biomass changes over time and across environmental conditions^8–11^. Thus, ATNs facilitate our understanding of how physiological responses to warming translate into species coexistence and biodiversity^7^. However, the ability of ATNs to derive sound predictions for large communities under changing environmental conditions has been challenged, stressing the need for more biological realism^6,12^. Indeed, a strong limitation of these models is that species are solely characterized by a set of biological rates that respond to temperature, such as metabolic or attack rates. Therefore, species are limited to physiological responses to warming, whereas the behavioural component is largely ignored. However, it is well established that species also respond to warming by changing their behaviour^13,14^. From a food web perspective, flexibility in species’ foraging behaviour is usually modelled using the principles of optimal foraging behaviour, which posit that consumers will modify their diet to maximize their energy intake. Under this hypothesis, models predict that behavioural changes help to support species coexistence in communities. However, it remains uncertain whether the premise of optimal foraging holds true in the context of rising temperatures and how behavioural changes prompted by warming translate into altered species coexistence. Therefore, integrating behavioural flexibility into food web models is critical to improve their accuracy in predicting the consequences of global warming^15–19^.

Energetic demands increase with temperature, but species can offset them by adopting various strategies to increase their energy intake. Species can actively forage on energetically more rewarding resources^13,20^, typically prey that are close to the maximum body mass that consumers can feed on^21^. Therefore, we expect that predators favour larger prey individuals (trait-based selectivity) at higher temperatures, reducing predator-prey body mass ratios (hypothesis H1a). Alternatively, individuals under high energetic stress may be driven by their increased demand for food and accept less rewarding (smaller), but more abundant prey upon random encounter (hypothesis H2a) leading to a reduced trait-based selectivity, and a trophic niche driven more by neutral processes (random encounter probability). These two hypotheses would lead to contrasting effects on communities^22^.

Trait-based selectivity may increase the interaction strengths between predators and larger prey, which could increase their extinction risk due to the lower abundance of large organisms^23^, therefore having negative consequences on species coexistence (hypothesis H1b). Alternatively, if neutral processes are found to be responsible for driving selectivity, predators would primarily forage on the most abundant species, which could result in a more even distribution of biomass across all prey size classes, enhancing the likelihood of species coexistence^18,24^ (hypothesis H2b). To test these hypotheses, we used a dataset of 2,487 stomach contents from 6 demersal fish species^25,26^ collected in the Baltic Sea between 1968 and 1978. This data set compiles body mass and information for the fish species, as well as body mass and biomass for the resource species observed in fish stomachs and in environment. We used this dataset to analyse the behavioural responses of fish to changes in temperature and resource availability (i.e., the total biomass of their prey). Subsequently, we addressed the consequences of these empirical relationships by integrating them into a food web model to predict how species coexistence changes with warming and the related foraging choices of consumers.

### Response to temperature and resource availability gradients

We used our database to document how consumer foraging behaviour responds to temperature and resource availability. The six fish species considered belong to two functional groups differing in body shape and foraging behaviour (flat, sit-and-wait predators versus fusiform, more active hunters). To describe the prey body mass distributions observed in fish stomachs (hereafter called the *realized distribution*) and in the environment (hereafter called the *environmental distribution*), we used empirical medians and standard deviations (Fig. 1). The environmental distribution defines what is expected if neutral processes drive fish diets: it represents the expected body mass distribution of the consumer diets if consumption was driven by density-based encounter rates only. However, the size distributions of prey in the environment and in consumer diets are usually not identical because consumers actively select prey individuals with specific body masses. We used the ratio of the realized and environmental distributions to calculate fish selectivity with respect to these different prey body masses to obtain a *preference distribution* (see conceptual explanation in Fig. SI 1), which describes consumer selectivity based on traits and consumer behavioural decisions (i.e. foraging behaviour). While traits define the fundamental trophic niche of a species (what a consumer can eat), behavioural decisions define the realised part of the fundamental niche. Therefore, a shift in behaviour does not necessarily imply a shift in the identity of prey species, but can simply lead to a shift in the individual traits that are selected, within or across different prey species. A more detailed description of the body mass relationships observed in consumer stomachs can be found in SI II.

**Figure 1:**
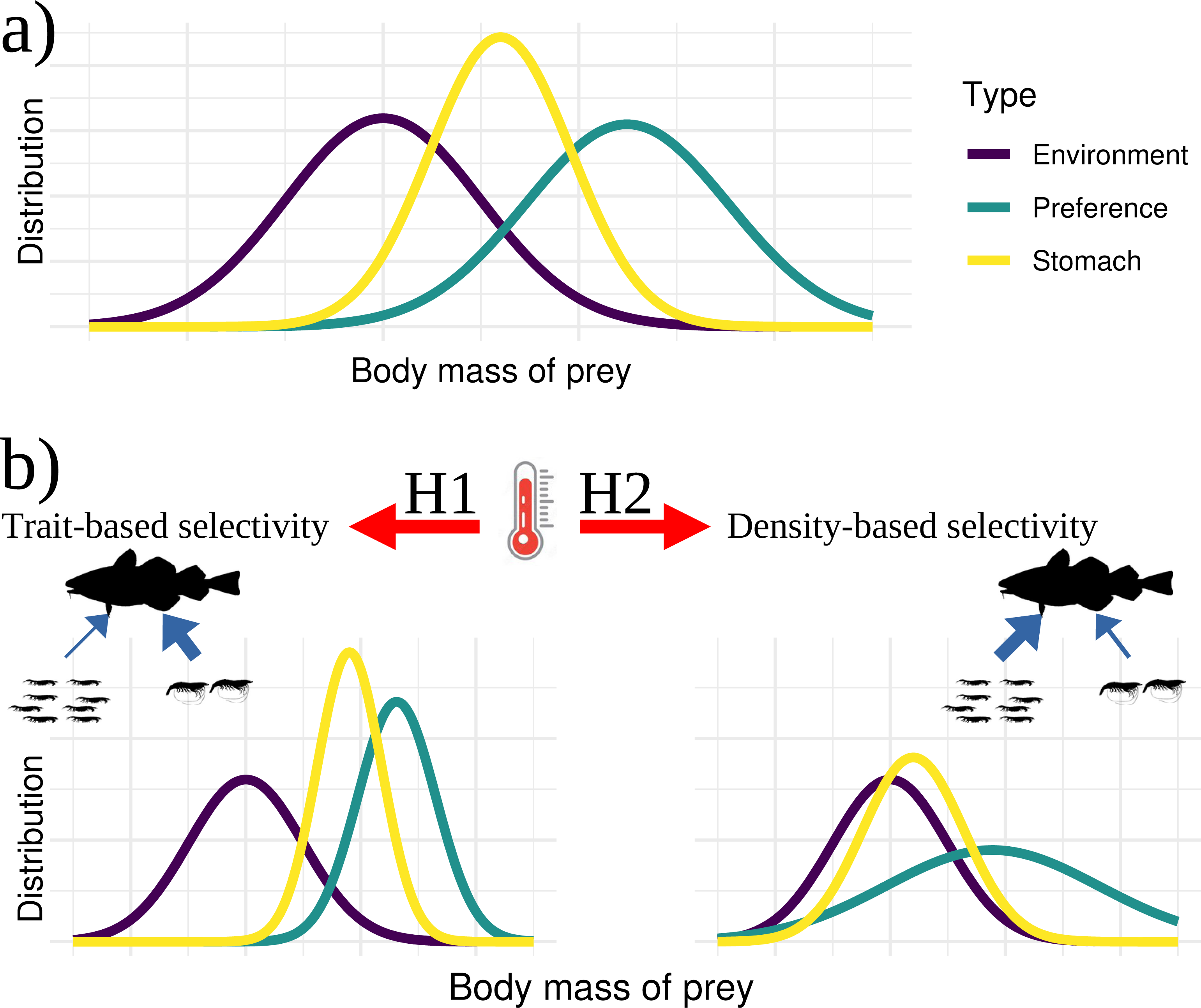
Conceptual representation of temperature impacts on the preference distribution. The preference distribution of prey body masses (turquoise) is estimated from the environmental (distribution of prey body masses in the environment, purple) and realised (distribution of prey body masses in consumers’ stomachs, yellow) body mass distributions (top panel). It represents how different prey body masses are selected by the consumers. Based on our two hypotheses, temperature increase can either: (H1, bottom left panel) lead consumers to preferentially select for larger species (trait-based selectivity), which can create an imbalance in the trophic fluxes (blue arrows): some species are preferentially selected even when their abundance is low, increasing their extinction risks and initiating large dynamical oscillations in species densities. Alternatively, temperature increase (H2, bottom right panel) leads consumers to have their diet more driven by encounter rates (density-based selectivity), which creates a stronger control of species with high biomass in comparison to less abundant, smaller ones, favouring coexistence of resource species and thus community species richness.

We tested how the median of the fish’s preference distributions was affected by the local resource availability and temperature conditions using a Bayesian linear model (note that we did not detect any strong correlation between resource availability and temperature; Spearman correlation = -0.167). The model included main effects for resource availability, temperature, fish functional group, and fish body mass, and the interactions between resource availability and temperature and between fish functional group and temperature. Including fish body mass as a covariate in the model ensures that our results are independent of predator size. We compared the full model to versions without the interaction between fish functional group and temperature and without fish functional group as a covariate using a “Leave One Out Cross Validation” approach (LOO-CV)^27^. The most parsimonious model used for the subsequent analyses was the one that did not include fish functional group or its interaction with temperature (results presented in SI III), indicating that the behavioural responses to temperature and resource availability were similar for fish species with different body shapes and foraging strategies. By contrast, we observed a significant interaction between resource availability and temperature on the median of the preference distribution (Table 1, Fig. 2) suggesting that behavioural responses of fish to temperature depend on environmental resource availability levels.

**Table 1:** Response of the median prey body mass to predator body mass and environmental gradients. Presentation of mean estimates and uncertainties (CI95), as well as effective sample size for the different predictors of the model. Rhat values were all lower than 1.003 for all estimated parameters. Total number of samples was 4000.

| Response of the median prey body mass of the preference distribution to |  |  |  |
| --- | --- | --- | --- |
| *Predictors* | *Estimates* | *CI (95%)* | *Effective samples* |
| Intercept | -1.06 | -3.14 – 1.09 | 1646 |
| Predator body mass | 0.55 | 0.41 – 0.70 | 2090 |
| Temperature | 0.18 | -0.04 – 0.38 | 2000 |
| Resource availability | 0.19 | -0.59 – 0.90 | 2017 |
| Temperature:<br>Resource availability | -0.07 | -0.14 – 0.01 | 1961 |
| Observations | 290 |  |  |
| R<sup>2</sup> Bayes | 0.279 |  |  |

**Figure. 2:**
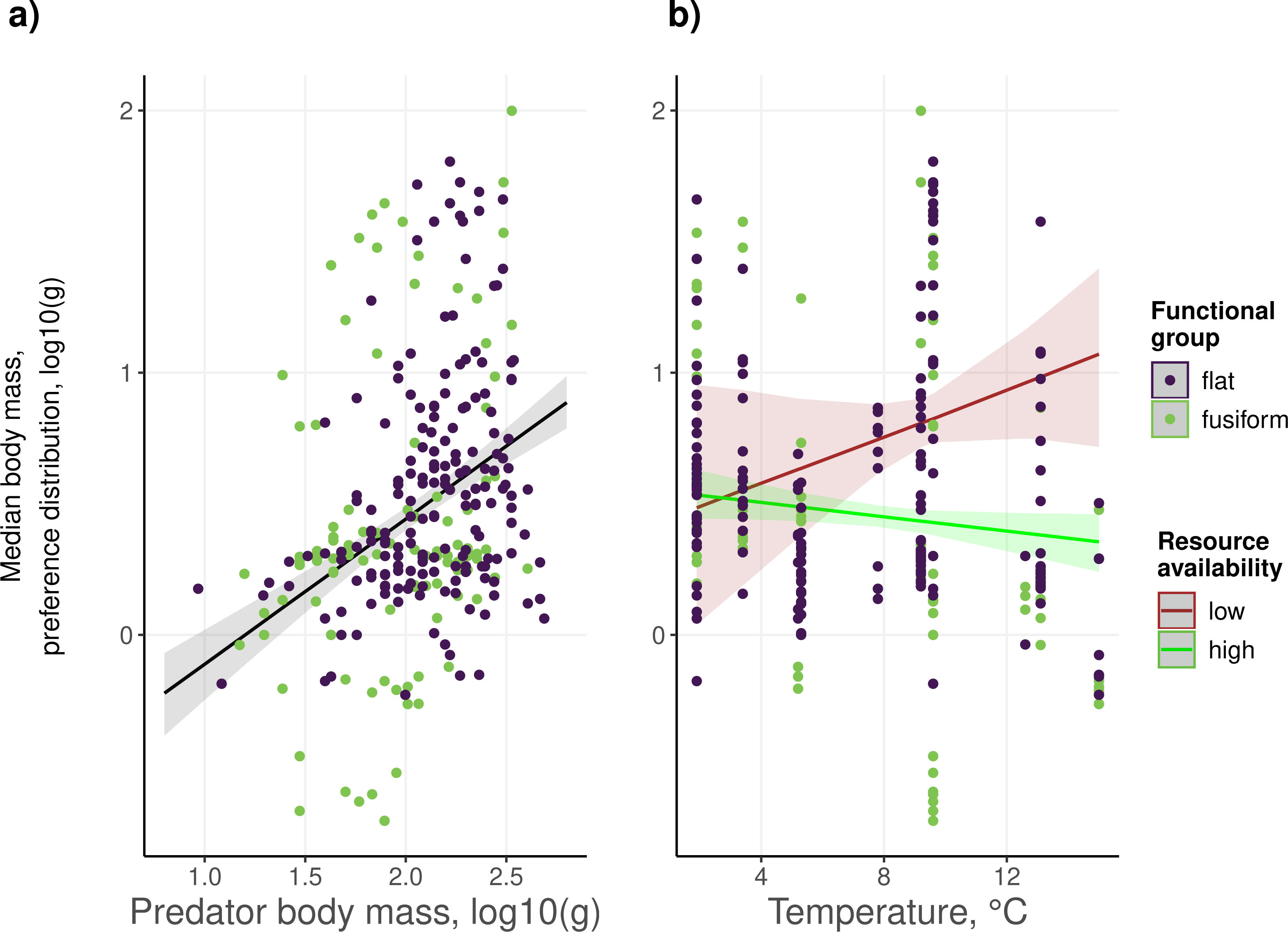
Response of the median prey body mass of the preference distribution. Effect of (a) predator body mass and (b) temperature and resource availability. Points represent log-transformed data across all resource availability levels and lines represent model predictions. Regression lines represent model predictions on the median of the preferred distribution when all other covariates are considered. The shaded areas show the 95% confidence interval on the predicted values. Low and high resource availability values correspond to the two modes of the bimodal distribution of resource availability values (presented in SI IV).

By examining the effect size and significance of the temperature effect along the resource availability gradient in our dataset, we found that the temperature effect is only significant at high levels of resource availability (Fig. 3). Here, fish tend to forage on smaller prey as temperature increases. This indicates that fish only change their feeding behaviour to temperature by foraging on smaller prey in warmer conditions (so supporting H2a against H1a) when resources are plentiful The energetic stress that warming imposes on individuals through increased metabolic rates should be mitigated by higher feeding rates at higher prey availability in more productive environments. Thus, because the effects of temperature and resource availability should cancel each other out, we expected a stronger behavioural response at low resource levels, where consumers must cope with maximum energetic stress (regardless of temperature). Under such stressful conditions, there may be no scope for predators to change their feeding behaviour as temperature increases, especially in the Baltic Sea where growth rates of fish species tend to be limited by resource availability in general^28,29^.

**Figure. 3:**
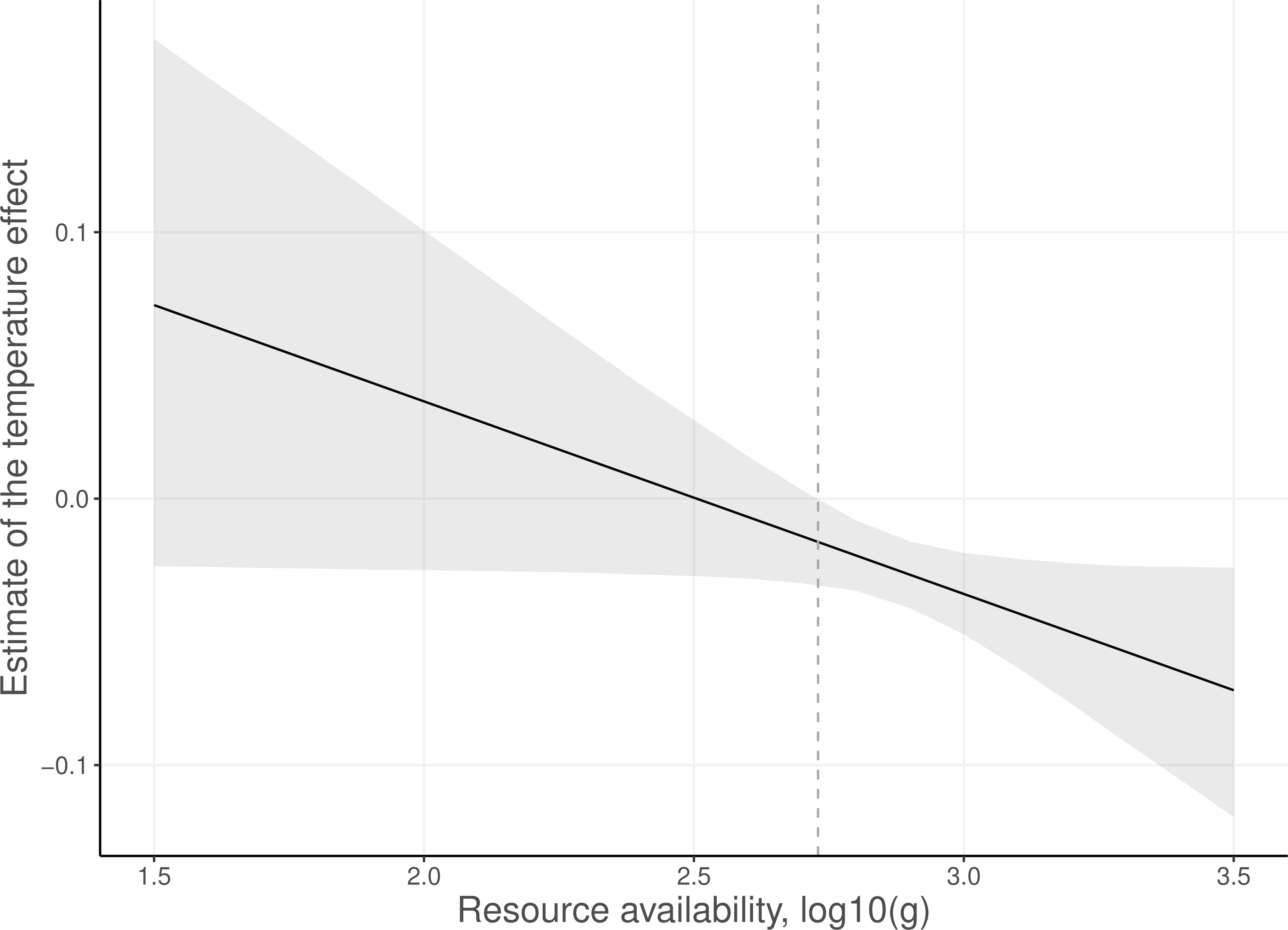
Effect of resource availability on the effect size of temperature on the median body mass of the preference distribution. The solid black line represents the effect size of the temperature effect calculated from the original model coefficients and the grey area shows the associated uncertainty (95% confidence interval). The vertical dashed line represents the resource availability threshold above which the temperature effect becomes significant (effect of 0 outside of the 95% confidence interval).

In more productive environments (See SI IV for a description of resource availability and of prey body mass structure), feeding behaviour may be less constrained, increasing the behavioural flexibility of the fish. We also observed that the width of the fish preference distribution decreases with temperature in the most productive environments (Table 2, Extended data 1), which means that this reduction in selected prey body masses is not explained by neutral processes but instead comes from fish actively foraging for smaller prey when temperature increases. This result challenges our hypothesis H2a, which suggested that foraging on smaller prey would be associated with a decrease in selectivity, but could be explained by the commonly observed type III functional response^23,30^. Here, when temperature increases consumers gradually shift their focus on more abundant resources (corresponding to smaller species) which comes at the cost of ignoring the less abundant, bigger ones, leading to a reduction in the width of the consumer trophic niche. With this behaviour, fish can satisfy their immediate demand for energy by targeting abundant species, but at the cost of missing out on larger and thus energetically more rewarding prey individuals, which can be critical to satisfy their energetic demands in the long term^31^. Indeed, fish metabolic rates increase with warming over large temperature gradients^32^ and do so faster than their feeding rates^33^, which can lead to the extinction of top predators due to starvation^34^. Combining this physiological starvation effect with our observed behavioural response indicates that consumption outside of the most efficient predator-prey body mass ratio should reduce energy flux through food webs, limiting the coexistence of consumer species^31,35^. The combination of physiological and behavioural warming effects could thus increase the likelihood of top predator extinction in food webs, which are usually considered key species for maintaining biodiversity and ecosystem functioning^36^.

**Table 2:**
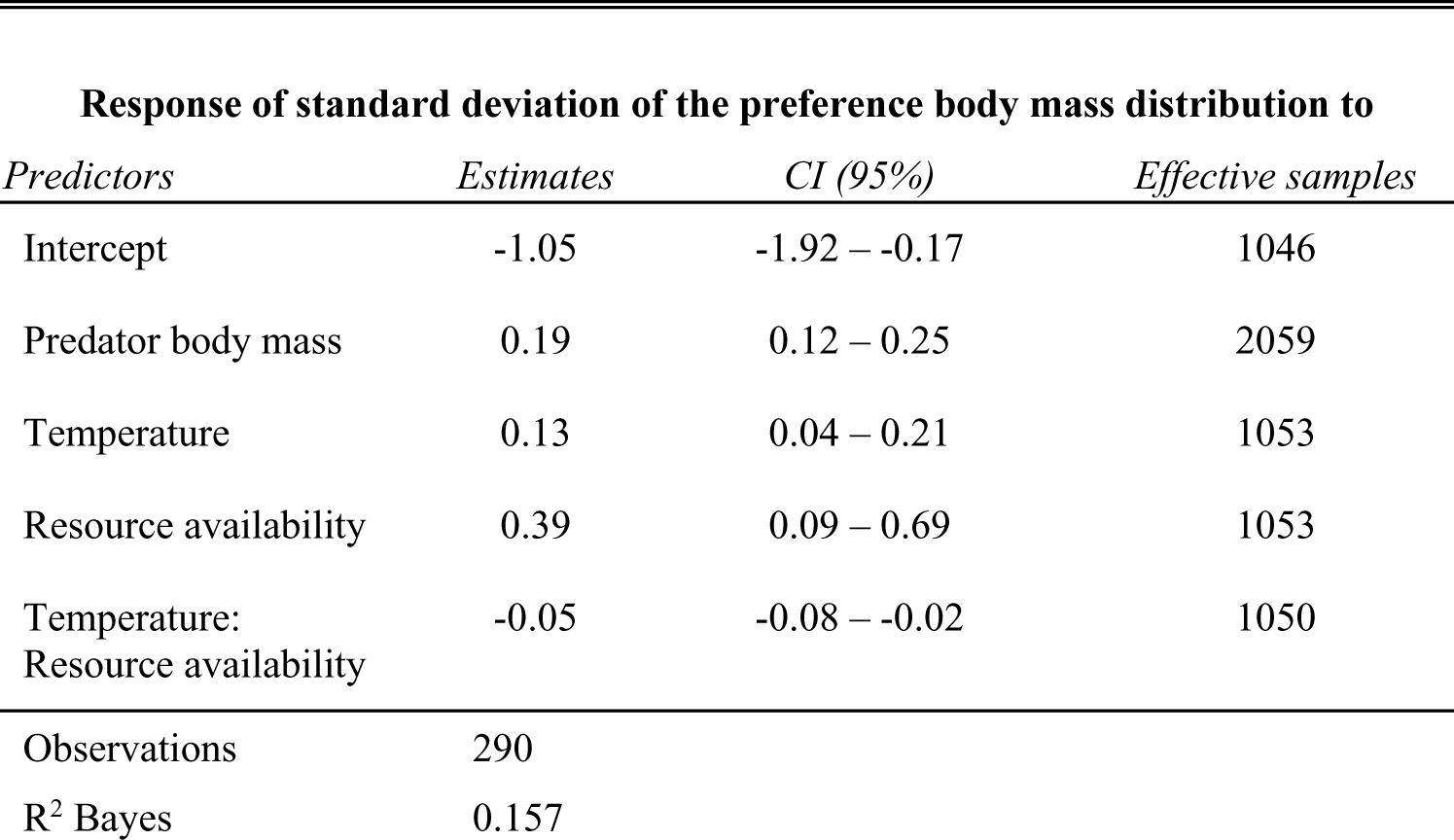
Response of the standard deviation of the body mass preference distribution to predator body mass and environmental gradients. Presentation of mean estimates and uncertainties (CI95), as well as effective sample size for the different predictors of the model. Rhat values were all lower than 1.002. Total number of samples was 4000.

### Consequences for species coexistence under global warming

Flexible foraging in response to varying local conditions is considered key to promote species coexistence^18,19,37^. The general assumption behind this notion is that consumer species will adapt their foraging strategies in order to maximize their energetic gains^38^. However, our results suggest that consumers tend to depart from this optimal foraging behaviour under stressful conditions. We explored the consequences of this behaviour using a dynamic population model, which predicts the temporal dynamics and coexistence of species in food webs (see Methods). We ran two versions of this model: one including changes of species diets based on local temperature and resource availability conditions as informed by our empirical results, and one model with static feeding links representing the classical food-web modelling approach. We simulated the dynamics for synthetic food webs of 50 species (30 consumers and 20 basal species) over a temperature gradient spanning from 0°C to 18°C to predict the number of species extinctions at different temperatures. Overall, we observed that models incorporating flexible foraging were more sensitive to warming, with more consumer extinctions over the temperature gradient (Fig. 4) and relatively few extinctions of basal resources (results presented in Fig. SI 6). These results were not affected by the functional response type, which is a free parameter in our modelling approach, but tended to weaken at very low levels of nutrient availability (i.e. productivity), consistent with our empirical results (model predictions for the different values are available in SI VI).

**Figure. 4:**
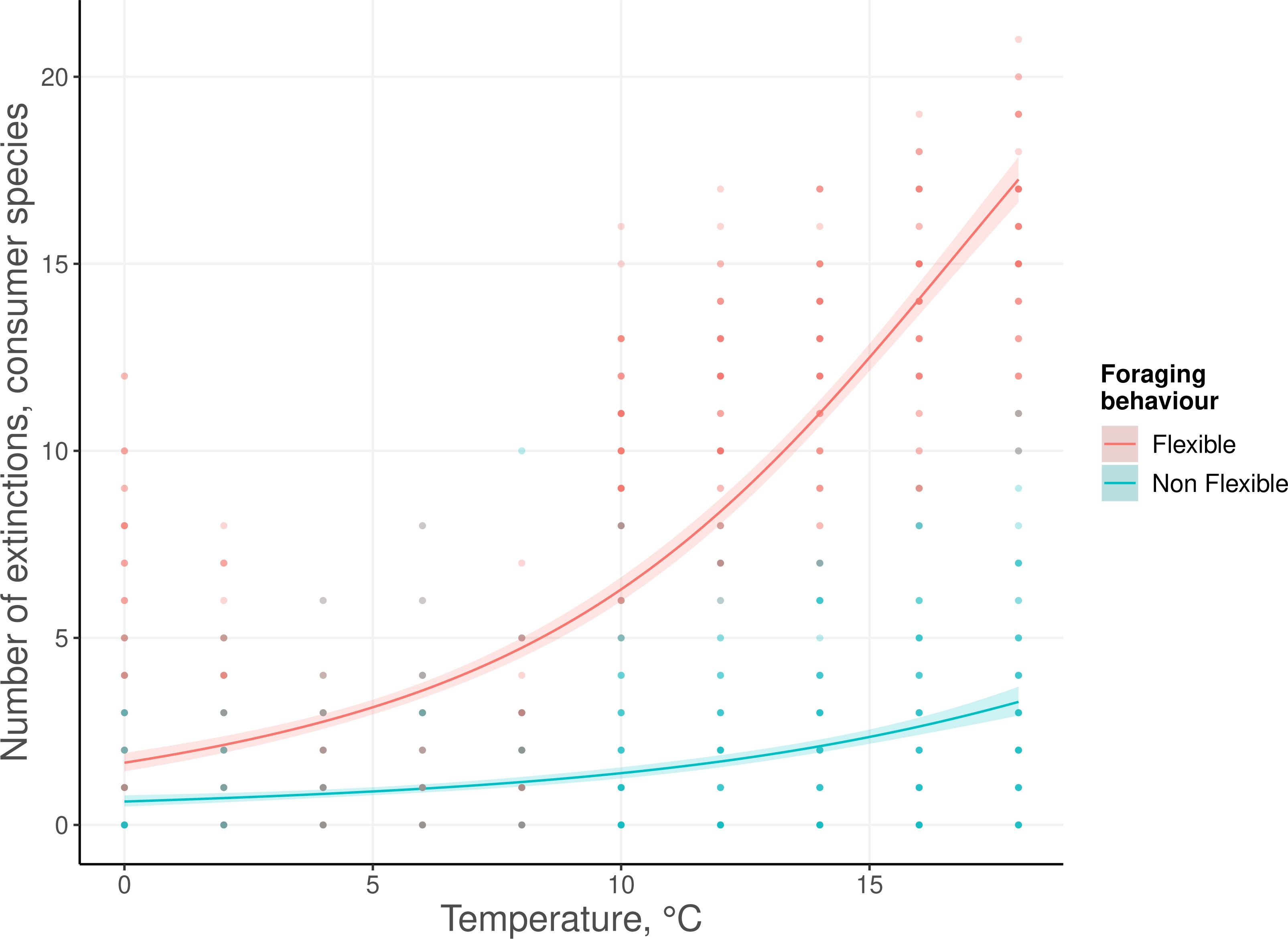
Number of consumer species extinctions predicted by the model at different temperatures (out of an initial richness of 30 consumers). Points represent the number of observed extinctions for each simulation. The blue line represents the model prediction on the average number of extinctions without the response of species’ foraging behaviour to local temperature and resource availability conditions considered, whilst the red line shows average predictions of extinctions when allowing for this flexibility. The shaded areas show the 95% confidence interval on the predicted values. Predictions were estimated using a GAM with a binomial link function.

Generally, food web models incorporating foraging behaviour are based on optimal foraging theory and thus lack a data-driven description of how the selectivity of consumer diets changes in a natural context. To address this gap, we developed a trait-based framework to document the response of foraging behaviour to temperature, which can be incorporated into predictive models of food web structure and species coexistence. Our approach can be generalized to other ecological variables that affect food webs and foraging behaviour, such as fear of predators or habitat complexity^39^. Finally, the effects documented here come from data sampled at rather low levels of temperature and resource availability. Therefore, it will be crucial to extend our regression models to warmer and more productive ecosystems to assess whether very high levels of productivity could balance the energetic stress related to warming, thus limiting behavioural responses in eutrophic environments.

A potential limitation of our approach is that the variations in temperature we employed were primarily driven by seasonal changes, which may not fully reflect the long-term dynamics associated with climate change as well as indirect effects through modification of habitat structure potentially interacting with the foraging behaviour of consumers or lowering oxygen concentrations. Similarly, response of social behaviour, like aggregation of predator or prey could influence the shape of the functional response but are rarely considered in the literature. Nevertheless, our hypotheses and interpretations are rooted in energetic budget calculations that stem from fundamental mechanistic principles concerning the impact of temperature and prey body masses on population growth rates. This mechanistic framework enabled us to formulate generalisable predictions regarding temperature effects across different ecological contexts. Consequently, we propose a testable explanation for the observed decline in biomass among the fish populations in the Baltic Sea^40^ . However, it remains possible that fish could exhibit increased fitness by consuming smaller, more abundant organisms under certain conditions. Hence, these predictions should now be confronted with a comprehensive examination of how the fitness of fish changes depending on temperature and behavioural choices, for example in more controlled experimental settings (e.g., mesocosms). Such experimental approaches could also consider other mechanisms leading to shifts in species diets and interactions in the context of temperature increase, e.g., ontogenetic shits^41^.

The effects of warming on the trait structure of communities and the distribution of trophic interactions^42^ are well documented, but a framework for integrating changes in feeding behaviour with a general modelling approach has been lacking. Closing this gap, our results stress the importance of accounting for foraging behaviour, to better understand and predict community responses to climate change, and challenge previous conclusions on this topic. Indeed, the discrepancies between the models with and without flexible foraging suggest that the classical approach, which only accounts for changes in species physiology^6,7^ completely ignoring behavioural aspects, may have overlooked a significant portion of community responses to warming. Importantly, our results show that, contrary to common expectation, behavioural responses to climatic stress can reduce the likelihood of species coexistence and thus decrease community biodiversity. The similarity in the responses of the two fish feeding strategies (sit-and-wait and active foraging) indicates some generality in our results, but it is important to investigate these patterns for a wider range of species and ecosystem types in future studies. For instance, consumer metabolic type has an important effect on the response of species to temperature^43^ and endotherms could respond differently than ectotherms such as fish.

## Conclusion

It is generally assumed that consumers respond to environmental conditions by making choices that maximize their energy intake^19,44^. This assumption has been used to derive several predictions in ecology about community structure and species coexistence, and is often considered a solution to May’s paradox^45^ that highly diverse communities are mostly mathematically infeasible, despite their widespread occurrence in nature. It is therefore commonly assumed that behaviour not included in these simple mathematical models is a strong driver of community organization and supports species coexistence^19^. We challenge this optimistic view of nature by demonstrating how consumer species can shift to less efficient foraging behaviour under stressful conditions, for instance when resources are scarce and when they face additional energetic stress due to warming. Therefore, the ecological conclusions built into the assumptions that flexible behaviour favours coexistence do not necessarily hold in the context of global warming. Our mechanistic modelling approach demonstrates the consequences of this observation, with more species extinctions in response to warming when flexible foraging is considered. This indicates that global warming may lead to a greater reduction in species coexistence than predicted by classical ecological models. Our findings thus challenge the general paradigm that flexible foraging should mitigate the consequences of global warming for natural ecosystems and call for a general data-driven theory approach to forecasting biodiversity and functioning in future ecosystems.

## Acknowledgements

We are profoundly grateful that Wolf E. Arntz collected and provided the valuable data set from his early work in Kiel Bay that we used in this study. We are also thankful to Astrid Jarre who digitized the stomach content data, Ute Jacob for her help in the early phase of this project and Carlos Melian for his friendly review of the manuscript. We gratefully acknowledge the support of: iDiv funded by the German Research Foundation (DFG–FZT 118, 202548816), BG, UB, BR, TB, MJ.

German Academic Exchange Service (DAAD, 57070483), GK.

European Research Council (ERC) under the European Union’s Horizon 2020 research and innovation programme (grant agreement no. 677232), MJ.

## Author contributions

BG, BR, UB conceptualised the study. BG, GK, BR, TB where involved in data compilation and code development and BG and BR analysed the data. BG wrote the first draft of the manuscript ad all authors contributed to its revision.

## Methods

### The Kiel Bay database

To estimate how species body mass preferences change depending on local environmental conditions we used the Kiel Bay database^26^. This publicly available database^25^ compiles the stomach contents of seven fish species (adult and sub-adult individuals) classified into two functional groups, based on their body shape and habitat use: fusiform and benthopelagic species (*Gadus morhua* and *Merlangius merlangus*) versus flat and demersal species (*Limanda*, *Pleuronectes platessa*, *Platichthys flesus*, and *Hippoglossoides platessoides*). This shape characteristic also corresponds to specific foraging behaviour^46^, where flat fish tend to be “sit-and-wait” predators, while fusiform fish are more actively foraging feeders. Together with taxonomic information, the database contains the body length of the fish species (rounded to the nearest integer in cm) as well as the abundance and biomass of the different prey species in fish stomachs and in their environments. Species in the environment were sampled with van Veen grabs directly along the trawling routes used for fish sampling. In total, 6 sampling sites (3 for each trawling route) were used for the bottom grab sampling, each site being sampled several times during the period covered by our dataset. To reduce biases associated with site specificity of benthos communities (i.e., sampled with the van Veen grabs), data from sampling sites were pooled together if they were harvested on the same date and corresponded to the same trawling route.

The dataset also contains sea temperature data (representing monthly average taken at 20m depth). For our analysis, we used a subset of the database for which it was possible to associate the content of fish stomachs to the environmental abundances of their prey (see the method section “*Filtering data and association between fish and environments”* for more details).

### Estimation of fish body masses

To allow for correspondences between our allometric food web model (that relies on species body masses for its parametrisation), the energetic approach discussed, and the empirical part of our study, we converted the body lengths of fish to body masses. We used a power law^47^: 𝐵𝑀_𝑖_ = 𝑎𝐵𝐿^𝑏^, were *BM_i_* and *BL_i_* represent the body mass and body length of fish *i*, respectively. *a* and *b* are constants that are species-and location-specific. Values for our parameters a and b obtained from fishBase^48^ are: *Gadus morhua*: a = 0.00708, b = 3.08; *Merlangius merlangus*: a = 0.00631, b = 3.05; *Limanda*: a = 0.00776, b = 3.08; *Pleuronectes platessa*; a = 0.00776, b = 3.06; *Platichthys flesus*: a = 0.00776, b = 3.07; *Hippoglossoides platessoides*: a = 0.00562, b = 3.09; *Enchelyopus cimbrius*: a = 0.00389, b = 3.08.

### Filtering data and association between fish and environments

To make comparisons between the distributions of prey observed in fish stomachs and the ones observed in the environment, we only used a subset of the Kiel Bay database for which we were able to associate information about a fish’s stomach contents to information about the related prey in the environment. We considered this association between fish and environment possible when they were sampled in the same area (i.e. bottom grabs carried out directly on the route of a bottom trawl) and within less than 31 days (see SI VII for a distribution of the differences). Then, associations were made between fish and environments from the same location that minimised the time difference between the fish and environmental sampling. This first filter reduced the number of fish used in our analysis to 2,487. From this subset, we pooled all individuals from the same fish species occurring at the same place on the same date (i.e., when they were harvested during the same sampling event) with the same body mass (in our case, body mass is a discrete variable as it was estimated from body length, rounded to the nearest integer in cm) into a unique entity for statistical analysis, which we hereafter call “statistical fish”. This choice is led by the allometric approach used in our analysis, where all individuals from the same species and with the same body mass are considered identical. This aggregation increases the quality of the estimation of the prey body mass distribution in stomachs at the cost of a lower statistical power for the analyses done on the shape of these distributions. For instance, with a high aggregation level, fewer data points are available to consider the effect of temperature on the average body mass of prey. This approach is therefore conservative as it reduces the probability of type 1 error. Lastly, to make sure that the sampling of the prey in the environment was representative of the statistical fish’s diet, we checked if the species composition in the environment matched that of the fish stomachs (<FUTURE reference to the data paper>). This led to the removal of 26 statistical fish where less than 90% of the biomass found in the diet corresponded to species also found in the environment. This resulted in a final dataset of 290 statistical fish, underpinned by 2,487 individuals. For our statistical analysis, we used fish body functional group as a covariate instead of fish species, as models based on fish functional group were always found to be more parsimonious (based on AIC).

### Fitting of gut content and environmental distributions

We used empirical medians and standard deviations to describe all environmental distributions of log_10_ body masses and realised distributions of each predator identity. Taxon-specific characteristics of the prey, such as body toughness, could bias the dietary distributions towards prey containing shells or skeletons. We assumed that prey with hard body parts are more likely to be detected in stomach contents than species composed of soft tissues (due to their higher digestion time) and weighted their occurrence by a correction factor of 0.8, (according to^49^) Overall, the trends and effects observed when including this correction were similar to those observed without correction, thus suggesting an absence of systematic biases (see SI VIII for an analysis without correction factor).

### Determining allometric species’ preferences

We assumed that a feeding event is defined by two independent probabilities: the probability for a consumer to encounter a prey of a certain body size *x* (defined by the environmental distribution *E*(*x*)) and the probability for a consumer to consume the prey when encountered (given by the preference distribution *P*(*x*)). Then, the realised distribution is proportional to their product:

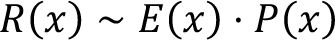

The preference distribution can therefore be expressed by the departure of the realised niche from the environmental distribution, or by filtering out the effect of species environmental availability from the realised distribution:

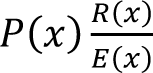

As such, the preference distribution is independent of seasonal changes in prey communities or any other factors that could lead to an altered size distribution of prey in the environment.

Theoretically, it is possible to compute continuous distributions *R* and *E* from observed body masses for the realised distribution, *r_i_* (*i*=1…*n*) and for the environmental distribution *e_i_* (*i*=1…*m*), respectively, with e.g. kernel density estimation, and compute:

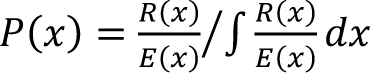

We chose, however, a more conservative approach that requires just a kernel density estimate for the environmental distribution *E*(*x*): Moments (i.e., mean, standard deviation, and skewness) of *P*(*x*) can be computed as weighted moments of the observed realised body masses *r_i_*with weights (i.e., the contribution of the datapoint to the estimation of the moments) *w_i_*=1/*E*(*r_i_*) being the inverse of environmental abundances. Thus, realised body masses that are highly abundant in the environment contribute less to the preference distribution, while those that are rare contribute more. Following^50^ and assuming 𝑊 = ∑_𝑖_ 𝑤, the mean *μ*, variance *σ*² and skewness *γ* of the preference distribution *P*(*x*) are:

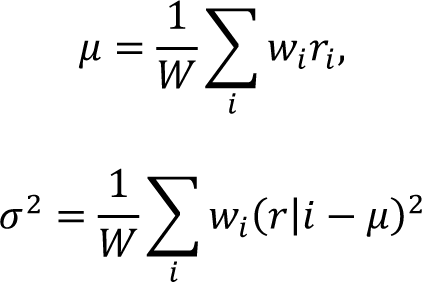

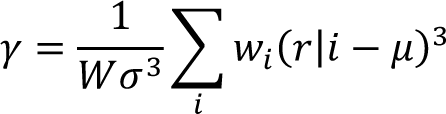

To assess changes in the distributions and how they depart from each other, we used variations in the point estimates (median and standard deviation).

### Statistical analyses

To fit the preference distributions, we used a Bayesian linear model to explicitly test our hypotheses. We started by predicting the median and standard deviation of the preference distributions using temperature, resource availability, fish functional group, and fish body mass as covariates, as well as interactions between temperature and functional group and temperature and resource availability.

Body masses used for the different distributions, as well as resource availability, were log-transformed for visualisation purposes and to meet homoscedasticity assumptions from linear models. The inclusion of fish body mass as a fixed effect in our model ensures that the other covariates are corrected by fish body mass and therefore independent of it. We first checked if fish functional group was an important predictor in our model using a “Leave-one-out” cross validation^27^, and finally simplified our model by removing fish functional group from the covariates (see SI III for a more comparison of the different models and for the presentation of posterior distributions). The log-transformation of the body mass and biomass variables were done to fulfil the assumptions of linear models.

### Dynamic model

To simulate the population dynamics, we used a previously published model^51^, based on the Yodzis and Innes framework^52^. The growth of consumer species *B_i_* is determined by the balance between its energetic income (predation) and its energetic losses (predation metabolism)

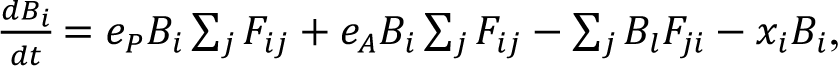

where *e_p_* = 0.545 and *e_a_* = 0.906 represent the assimilation efficiency of a consumer foraging on plants and animals, respectively^53^. *x_i_* defines the metabolic rate of species *i*, which scales allometrically with body mass:

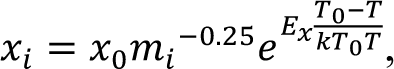

where *x_0_ = 0.314* is the normalisation constant^51^, *E_x_* = -0.69 is the activation energy of metabolic rate^7^, *k* the Boltzmann constant, *T_0_ = 293.15* the reference temperature in Kelvin and *T* the temperature at which the simulation is performed. The trophic interactions are determined using a functional response *F_ij_*that describes the feeding rate of consumer *i* over resource *j*:

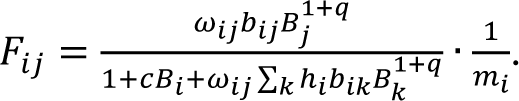

*b_ij_* represent the species-specific capture and is determined by predator and prey body masses: *b_ij_* = *P_ij_ L_ij_*.

It corresponds to the product of encounter probabilities *P_ij_* by the probability that an encounter leads to a realised predation event *L_ij_*. As such, the parameters encode neutral processes (encounter probabilities) and trait-based selectivity, as the distribution *L_ij_* represents the fundamental trophic niche of consumer *i*, i.e. the set of prey it can consume based on its traits. Both quantities are determined by species body masses. We assume that encounter probability is more likely for species with higher movement speeds of both consumer and resource species:

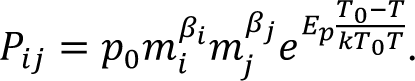

Since movement speed scales allometrically with consumer and resource masses^54^, we drew *β_i_*and *β_j_* from normal distributions (consumer: *μ*_β_ = 0.47, *σ*_β_ = 0.04, resource: *μ*_β_ = 0.15, *σ*_β_ = 0.03), following^51^. Activation energy *E_p_*is equal to -0.38, from^7^. *L_ij_*is assumed to follow a Ricker curve^51^, defined as:

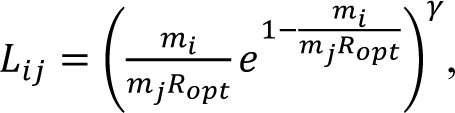

where the optimal consumer-resource body mass ratio *R_opt_ = 71.68* was calculated from the observed realised interactions in our dataset. We used a threshold *L_ij_ < 0.01* under which values were set to 0, assuming that consumers do not consider prey which are too small or too large. This fundamental niche described by *L_ij_* was used to generate our food webs, as we considered that species *i* consumes species j when *L_ij_* >0.01. The handling time *h_ij_*of *i* on *j* is defined as:

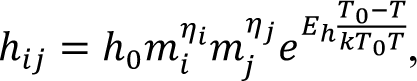

where the scaling constant *h*_0_ was set to 0.4 and the allometric coefficients for 𝜂_𝑖_ and 𝜂_𝑗_ were drawn from a normal distribution with mean and standard deviation of -0.48 and 0.03 for 𝜂_𝑖_ and of -0.66 and 0.02 for 𝜂_𝑗_. *E_h_* is equal to 0.26. *c_i_* represents the species-specific interference competition defined as:

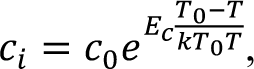

Where the scaling constant *c*_0_ was drawn from a normal distribution of mean 0.8 and standard deviation 0.2. The activation energy *E_c_* is equal to -0.65. The hill exponent *q* is a free parameter coding for the type of functional response and is set to 1.2 by default. The term *w_ij_* informs on species selectivity^55^, describing the foraging effort of a given consumer on part of its fundamental niche (based on traits only, as described by the *L_ij_*). For the models without behavioural expectations, we defined *w_ij_* for every resource *j* as 1 over the number of prey species of consumer *i*. This parametrisation, which corresponds to what is usually done in modelling studies, means that a consumer will split its foraging effort equally among its prey, independent of the local environmental conditions. When flexible behaviour was included in the model, the values of *w_ij_* were determined by the empirical preference distributions associated with fish. Our empirical distributions were characterised by the 3 first moments (mean, standard deviation, and skewness) which can be used to estimate the parameters of a skewed normal distribution, named location (ξ), scale (ꞷ), and shape (α), respectively. As for the mean and standard deviations, we used linear models to relate these parameters to consumer body mass, as well as the interaction between resource availability and temperature. The scale parameter (ꞷ) is constrained to positive values, so we used a generalised linear model with a log link function for this specific parameter. For each consumer, based on its body mass, as well as local temperature and resource availability conditions, we predicted the complete parametrisation of a skewed normal distribution. The preference of consumer *i* on resource *j* was estimated with the values returned by the probability density functions associated to each parameterisation for the different resource body masses.

To maintain the comparability with the model without flexible behaviour, the *w_ij_* values were transformed so that their sum was equal to 1 for each consumer. The biomass dynamic of the basal species *i* is defined as:

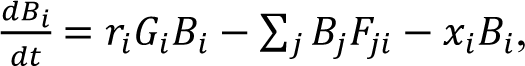

where 𝑟_𝑖_ = 𝑚^−0.25^ defines the species growth rate (𝐸_𝑟_ is the associated activation energy, set to 0.25). *G_i_* is the species-specific growth factor, determined by the concentration of two nutrients *N_1_* and *N_2_*:

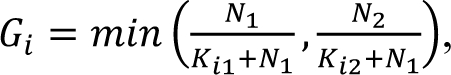

Where *K_il_* determines the half saturation density of plant *i* nutrient uptake rate, determined randomly from a uniform distribution in [0.1, 0.2]. The dynamic of the nutrient concentrations is defined by:

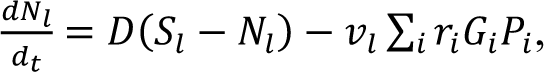

Where *D = 0.25* determines the nutrient turnover rate and *S_l_ = 5* determines the maximal nutrient level. The loss of a specific nutrient *N_l_* is limited by its relative content in plant species biomass (*v*_1_=1, *v*_2_=0.5). We ran our model on food webs of 50 species, composed of 30 consumers and 20 basal species. A link was drawn between two species *i* and *j* when *L_ij_ > 0*. For each temperature we ran 50 replicates of the two versions of the model (with and without flexible behaviour) using R 4.0.0-2 and with an updated version of the ATNr package^56^,using the “Unscaled ATN with nutrients” version. The parameters used correspond to the one provided by the package and summarised in SI IX. We fitted a GAM model on this number of extinctions. The code used for our analysis is available at^57^ https://zenodo.org/doi/10.5281/zenodo.10554928

## Data Availability

The data used for this study can be accessed at: https://doi.org/10.25829/idiv.3547-rtgq13 (see^25^)

## Code Availability

The code used for this study is available at: https://zenodo.org/doi/10.5281/zenodo.10554928 (see^57^)

