## Supplementary Information for "Flexible foraging behaviour increases predator vulnerability to climate change"

**Supplementary information I: Relations between environmental, realised and preference distributions.**

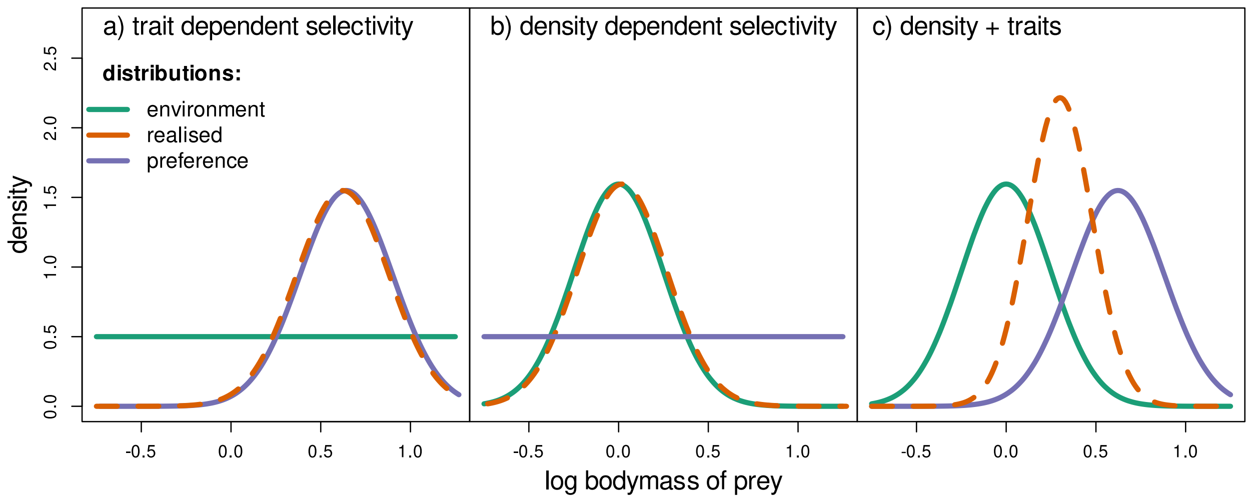

**Figure SI 1:** Illustration of the different fish prey body mass distributions. The environmental distribution (green) represents the distribution of prey body mass in the ecosystem, the realized distribution (dashed red) represents the body mass of the prey in a consumer stomach, and the preference distribution (blue) represents the selectivity of a consumer for a specific prey body mass. a) All of the log prey body masses are equally represented in the environment, so the distribution of prey body masses observed in a consumer's gut represents the body masses on which it actively foraged (its preference distribution) and predation is driven by trait selectivity only (hypothesis 1). b) The body mass distribution of the prey observed in the gut and in the environment are equivalent, so the prey consumed by the predator were entirely driven by encounter probabilities (i.e. a neutral process), implying no active selectivity over specific prey size classes (hypothesis 2). Panels a) and b) represent extreme scenarios while real-world data are more likely to be described by two different distributions, as in c) where the body mass distribution of prey observed in the stomach and in the environment differs, so that the consumer specifically forages on some prey body masses that are represented by the preference distribution. High values in the preference distribution represent body masses that are over-represented in fish stomachs compared to what is available in the environment.

### Supplementary information II: Response of the realised body mass distribution to predator body mass and environmental distributions

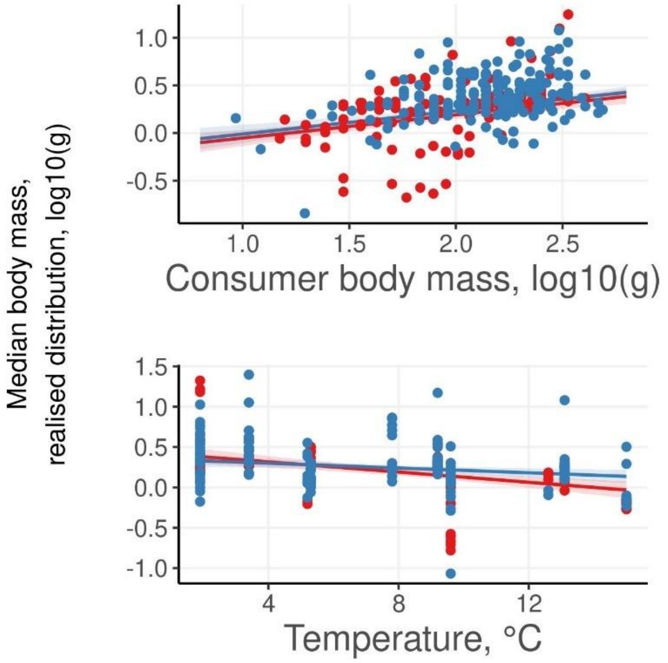

**Figure SI 2:** Response of the median body mass of the realised prey body mass distribution. Response to predator body mass (a), and temperature (b) for the two fish functional groups. Points represent non-transformed data across all productivity levels and lines present model predictions. Regression lines represent model's predictions for the median body mass of the realised distribution when all other covariates are considered. The shaded areas show the 95% confidence interval on the predicted values.

**Table SI 1:** Model coefficients. The model predicts the response of the median body mass of the realised distribution to predator traits and environmental gradients. Rhat values were all lower than 1.002. Number of samples was 4000.

| Response of the median prey body mass of the realised distribution to | | | |
|---|---|---|---|
| ***Predictors*** | ***Estimates*** | ***CI (95%)*** | ***Effective samples*** |
| Intercept | -0.95723 | -2.2867 – 0.3846 | 1809 |
| Predator body mass | 0.241989 | 0.1441 – 0.3399 | 3550 |
| Temperature | 0.036672 | -0.0932 – 0.1734 | 1759 |
| Productivity | 0.305422 | -0.1463 – 0.7688 | 1763 |
| shapefusiform | 0.083483 | -0.0592 – 0.2259 | 2521 |
| temperature:shapefusiform | -0.01697 | -0.0333 – -0.0003 | 2406 |
| temperature: resource availability | -0.0192 | -0.0672 – 0.0263 | 1755 |
| Observations | 290 | | |
| $R^2$ Bayes | 0.287 | | |

### Supplementary information III: Comparisons of model performances and diagnostics

**Table SI 3**: comparisons of model performences. The first row corresponds to the complete model, including all pairwise interactions (pred.BM:(resource availability), pred.BM:temperature, temperature:(resource availability) and temperature:shapefusiform). The second row corresponds to the model without the interaction between temperature and fish functional group. The third row corresponds to the model without fish functional group as a covariate. Δ elpd stands for differences in expected log pointwise predictive density (representing the goodness of model predictions) and Δ se its standard deviation (representing the uncertainty in the observed differences)

| | Δ elpd | Δ se |
|---|---|---|
| Complete model | 0.0 | 0.0 |
| -(functional group) x temperature | -4.9 | 3.8 |
| -functional group | -6.1 | 4.9 |

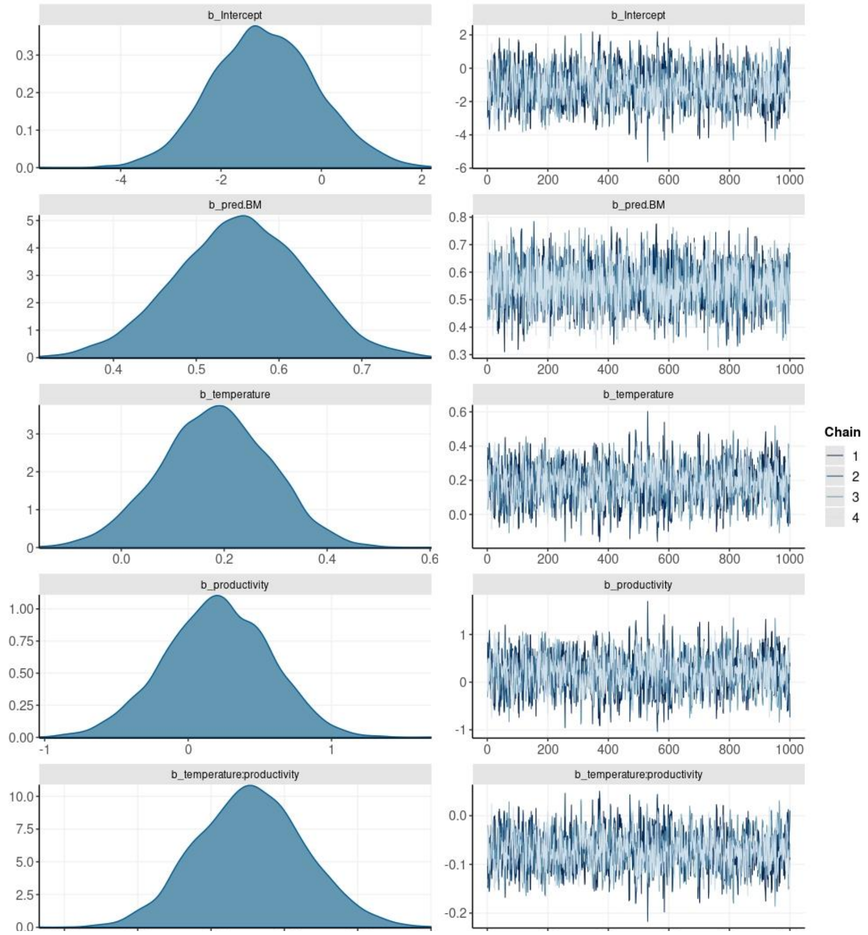

**Figure SI 3:** posterior distributions and chain mixing of the model without functional groups as a covariate

**Supplementary information IV: Environmental characteristics**

Overall, the different environments considered were characterised by two contrasted levels of productivity, leading to a bimodal distribution.

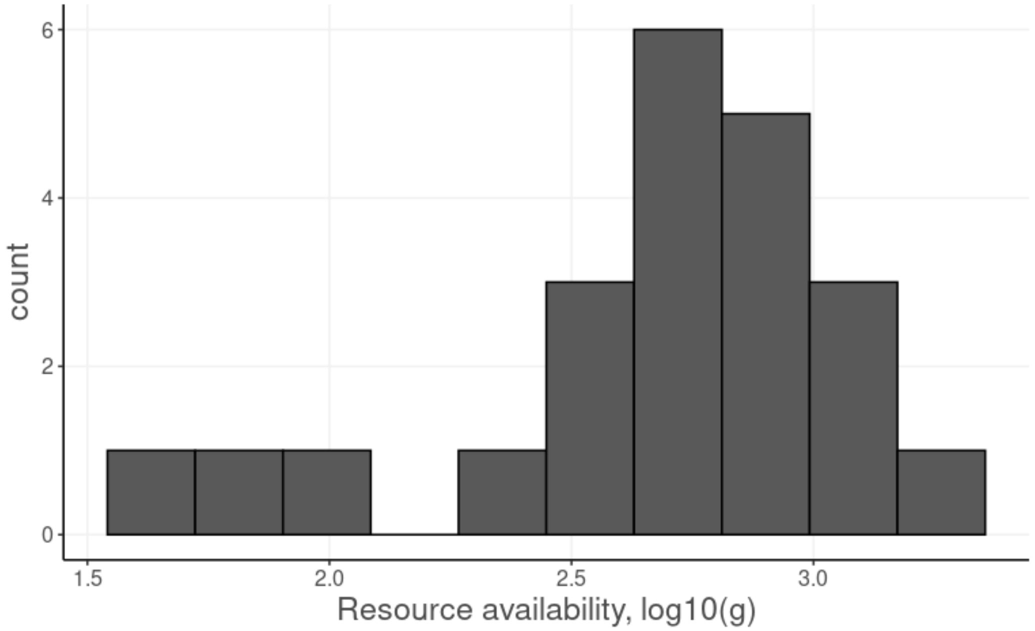

**Figure SI 4:** distribution of the resource availability for the different environments. Values are log10 transformed from grams (n = 22)

Associated to these differences, we observed that the body mass distribution of the prey species (median and standard deviation) was responding differently to temperature depending on productivity values (Figure SI II):

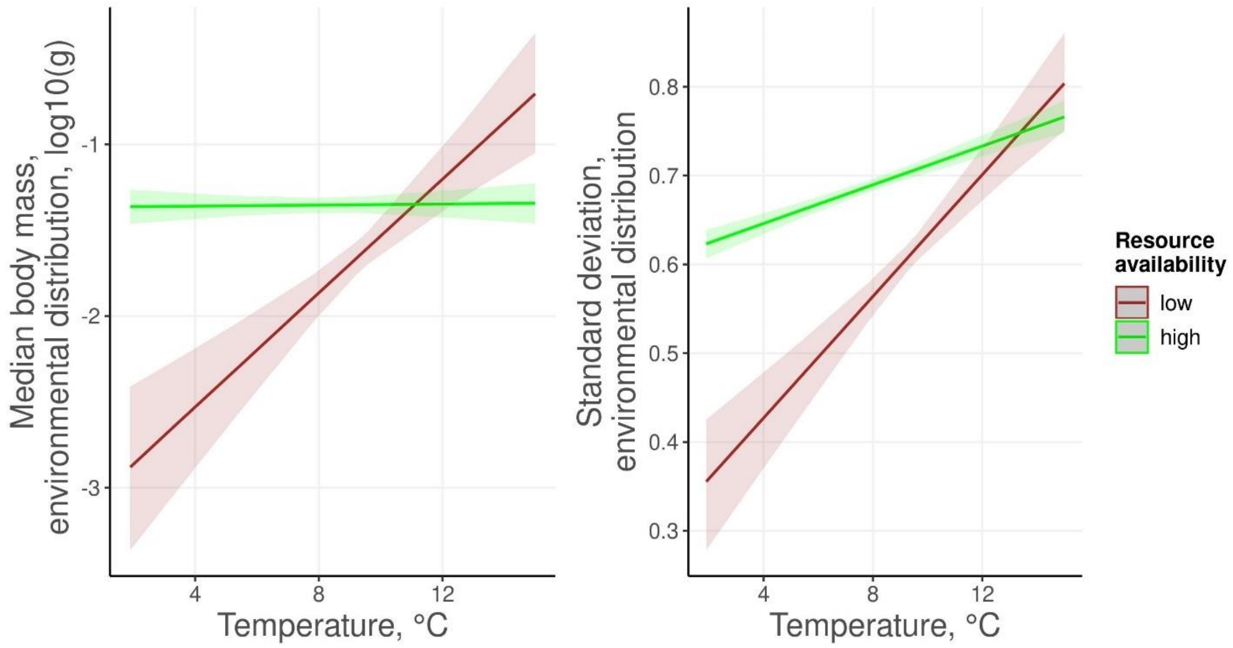

**Figure SI 5:** response of the body mass structure of the resource species (environmental distributions) to temperature and resource availability. Lines represents the model predictions and shaded areas their 95% confidence intervals

**Table SI 4:** model estimate for the prediction of median and standard deviation of the environment distributions. All Rhat values were lower than 1.006 indicating good chain mixing.

| | **Median of BM** | | **Standard deviation of BM** | |
|---|---|---|---|---|
| *Predictors* | *Estimates* | *CI (95%)* | *Estimates* | *CI (95%)* |
| Intercept | -7.54 | -9.76 – -5.44 | -0.45 | -0.80 – -0.13 |
| Temperature | 0.56 | 0.35 – 0.78 | 0.09 | 0.06 – 0.13 |
| Resource availability | 2.29 | 1.55 – 3.08 | 0.39 | 0.28 – 0.51 |
| Temperature: Resource availability | -0.21 | -0.28 – -0.13 | -0.03 | -0.04 – -0.02 |
| Observations | 290 | | 290 | |
| $R^2$ Bayes | 0.212 | | 0.311 | |

### Supplementary information V: Effect of temperature on the extinctions of basal species

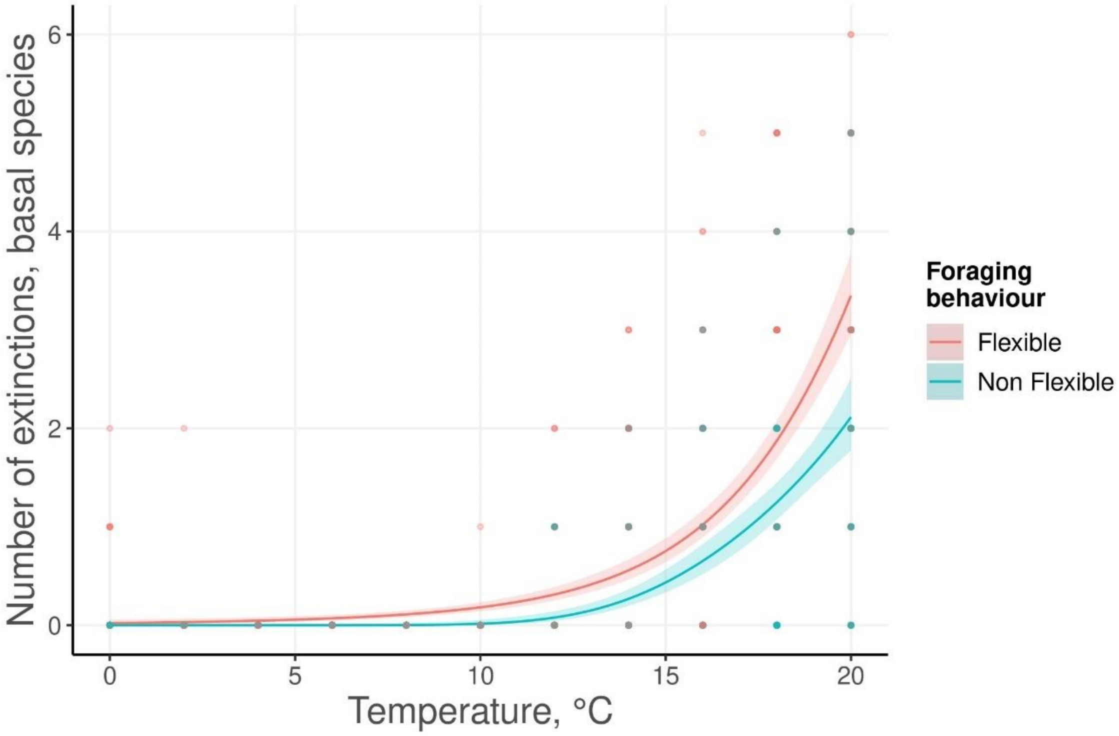

**Figure SI 6:** Number of basal species extinctions predicted by the model at different temperatures (out of an initial richness of 20 basal species). Points represent the number of observed extinctions for each simulation. The blue line represents the model output with adaptation of species diets to local temperature and productivity conditions considered, whilst the red line shows extinctions without allowing for this adaptation. The shaded areas show the 95% confidence interval on the predicted values. Predictions were estimated using a GAM with a binomial link function.

**Supplementary information VI: Effect of nutrient availability and predators' functional responses type on predictions about species coexistence.**

As maximum nutrient availability (variable $S_i$) and shape of the functional response ($q$) are not empirically informed, we analysed how sensitive to these two parameters model's predictions are. We varied $S_i$ from 5 to 40 and q from 1 to 1.5 (Fig. SI 7). We can here observe that few differences occur between our two models at very low productivity levels while more extinctions occur when simulating communities with models incorporating flexible foraging behaviour. This is consistent with our experimental results showing an effect of temperature on consumers' diet at higher productivity levels only. Consistently to what is expected (Binzer et al. 2012, Binzer et al. 2016), higher productivity values balance the effect of temperature increase when foraging behaviour is not considered. However, when we incorporated the foraging behaviour described in our experimental results, we can see that this compensatory effect disappears. At higher productivity values we can see a more complex response from model incorporating foraging behaviour. This increase in number of extinctions observed when temperatures becomes very low could correspond to unstable dynamics explained by a relatively high inflow of energy to species from the higher trophic levels while their metabolic rates remains low, consistently with Gauzens et al. 2020 The type of the functional response used resulted in more variations on the number of extinctions observed, with differences in extinctions between the two different models increasing with q (Fig. SI. 8).

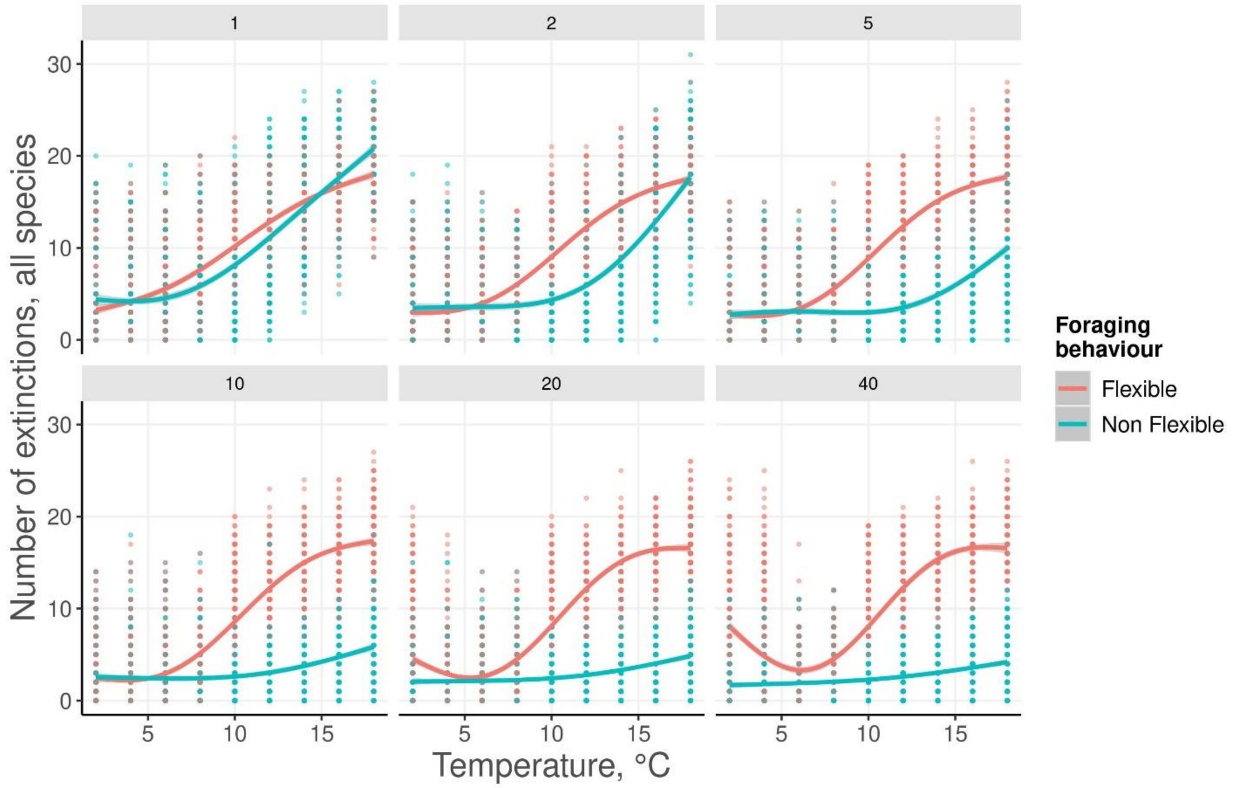

**Figure SI 7:** Effect of different levels of nutrient availability (variable S, values on top of the panels) on the number of extinctions predicted by the model. Simulations where ran with all hill exponent (q) values. Points represent the number of observed extinctions for each simulation. The blue line represents the model output with adaptation of species diets to local temperature and productivity conditions considered, whilst the red line shows extinctions without allowing for this adaptation. The shaded areas show the 95% confidence interval on the predicted values. Predictions were estimated using a GAM with a binomial link function.

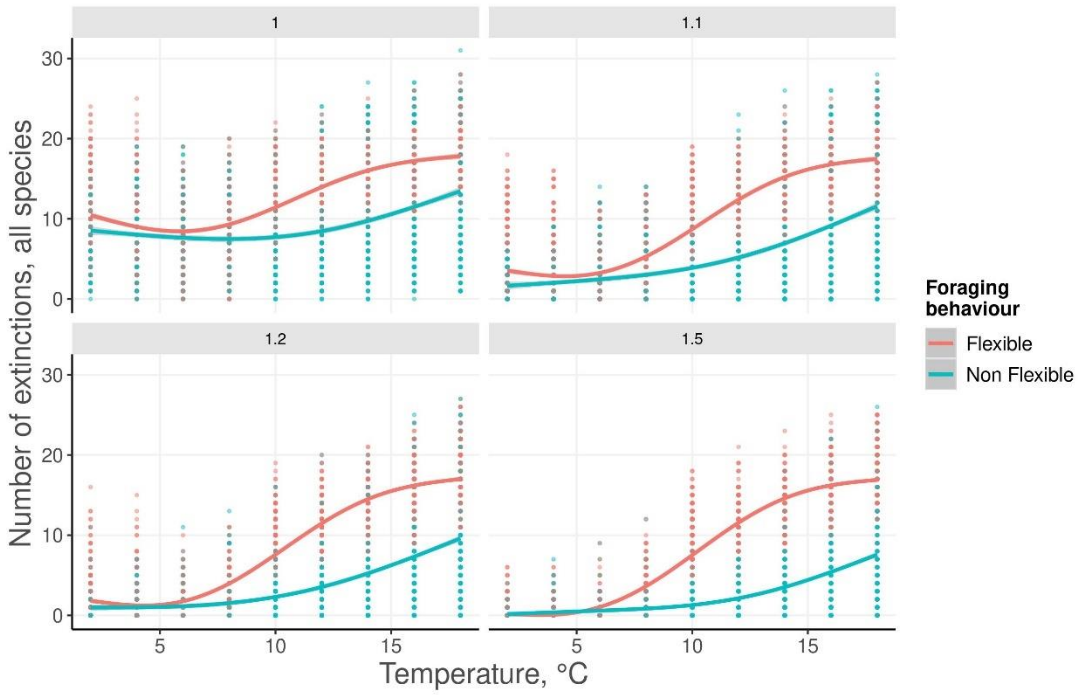

**Figure SI 8:** effect of the choice of functional response type (parameter q, values on top of the panels) on the number of extinctions predicted by the model. Simulations where ran for all level of maximum nutrient concentration (S). Points represent the number of observed extinctions for each simulation. The blue line represents the model output with adaptation of species diets to local temperature and productivity conditions considered, whilst the red line shows extinctions without allowing for this adaptation. The shaded areas show the 95% confidence interval on the predicted values. Predictions were estimated using a GAM with a binomial link function.

### Supplementary information VII: Sampling output and fish environments associations

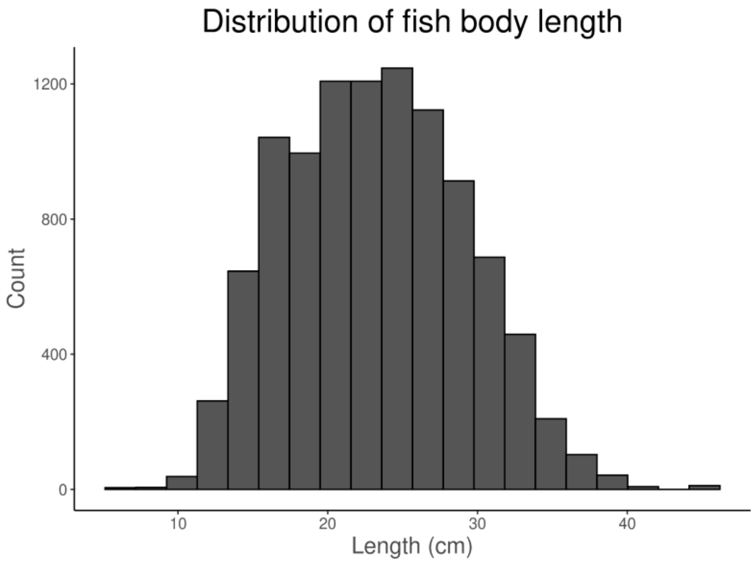

**Figure SI VII.1:** Distribution of fish body lengths (cm) in our database.

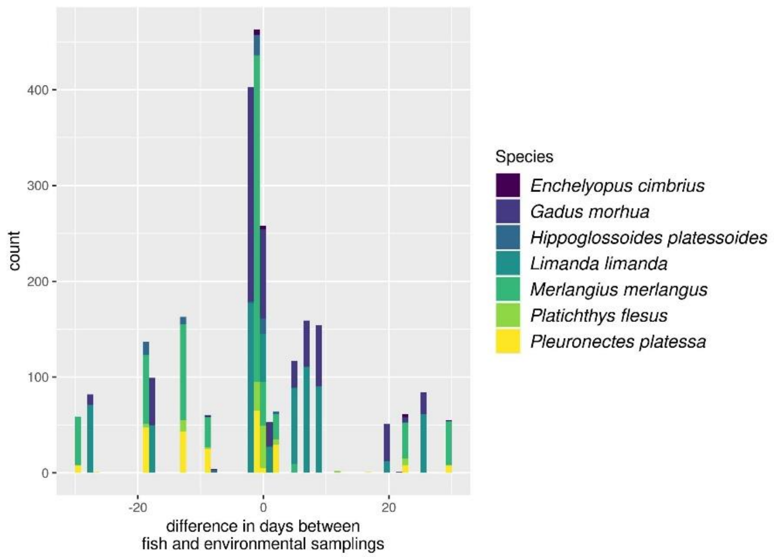

**Figure SI VII.2:** Distribution of temporal distance (days) between fish sampling and associated benthic samplings

**Supplementary information VIII: Effect of considering different detection probabilities for prey in stomachs**

As prey composed of soft tissues only are supposed to be less likely to be detected because of a faster digestion time, we corrected our observation by multiplying the abundance of species with hard body parts by 0.8. This was done to mirror the importance of these species that should persist longer in stomachs. As we are missing a general framework to properly describe how digestion time changes for the different species we used a unique correction factor that is a free parameter in our model (prey are either easy or difficult to digest, Table SI XI.2). We here present the results without this correction factor. Overall, these results remain qualitatively similar to the ones presented in the main text.

**Figure SI VIII.1:** Response of the median prey body mass of the preference distribution. Effect of (a), temperature (b) at different productivity levels. Preference values are based on data not corrected by prey digestibility. Points represent non-transformed data and lines present model predictions. The shaded areas show the 95% confidence interval on the predicted values.

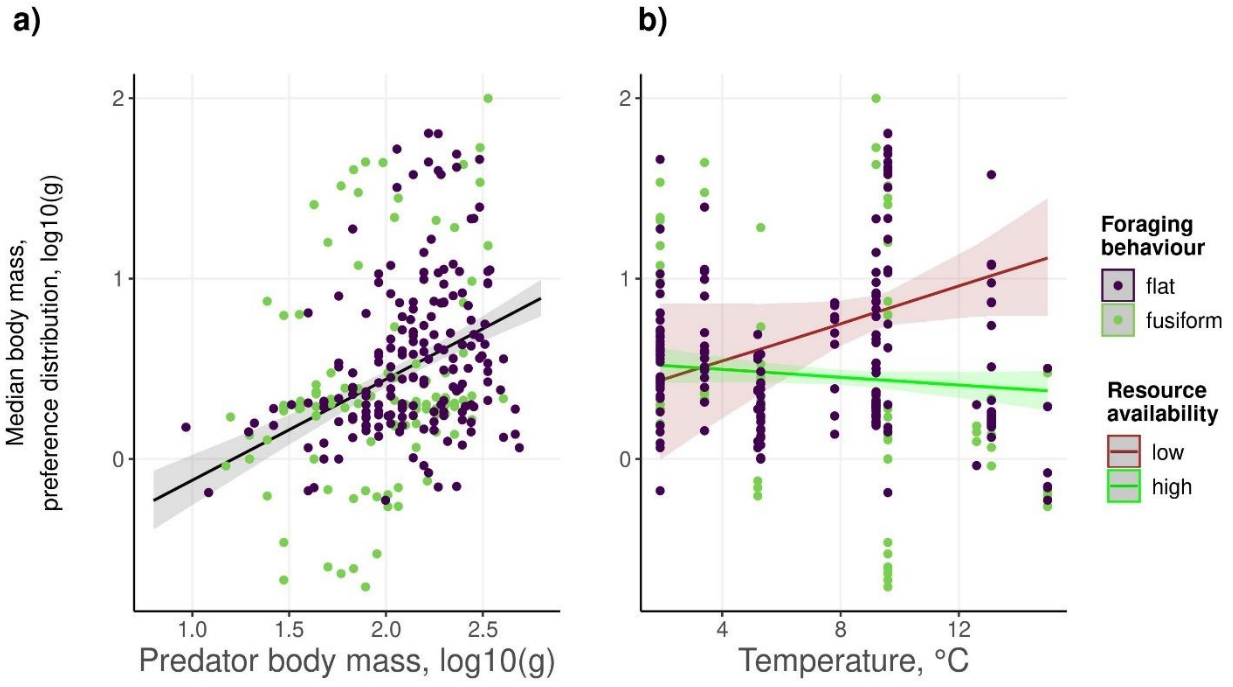

**Table SI VIII.1:** response of the realised distribution to predator body mass and environmental gradients when preferences are estimated with realised distributions non-corrected by prey digestibility. Rhat values were all lower than 1.005. Number of samples was 4000.

| | **Response of the median prey body mass of the preference distribution to** | | |
|---|---|---|---|
| *Predictors* | *Estimates* | *CI (95%)* | *Effective Samples* |
| Intercept | -1.32 | -3.35 – 0.63 | 2071 |
| Predator body mass | 0.56 | 0.41 – 0.72 | 2252 |
| Temperature | 0.2 | -0.0 – 0.41 | 1685 |
| Resource availability | 0.26 | -0.43 – 0.97 | 1805 |
| Temperature: Resource availability | -0.08 | -0.15 – -0.01 | 1754 |
| Observations | 290 | | |
| $R^2$ Bayes | 0.279 | | |

We can observe that the absence of correction factor does not qualitatively change the trends observed for the preference distributions. We can only detect slight changes in the model estimates.

**Table SI VIII.2:** Classification of species' digestibility Classification of species' digestibility

| Prey species | Class | Digestibility |
|---|---|---|
| *Abra alba* | Bivalvia | Hard |
| *Aloidis gibba* | Bivalvia | Hard |
| *Amphicteis gunneri* | Polychaeta | Easy |
| *Amphipoda spp.* | Malacostraca | Easy |
| *Anaitides spp.* | Polychaeta | Easy |
| *Anthozoa spp.* | Anthozoa | Easy |
| *Aphia minuta* | Actinopterygii | Hard |
| *Aphroditidae spp.* | Polychaeta | Easy |
| *Arenicola marina* | Polychaeta | Easy |
| *Ascidiacea spp.* | Ascidiacea | Easy |
| *Astarte spp.* | Bivalvia | Hard |
| *Balanus spp.* | Hexanauplia | Hard |
| *Brada villosa* | Polychaeta | Easy |
| *Capitella capitata* | Polychaeta | Easy |
| *Carcinus maenas* | Malacostraca | Hard |
| *Cardium fasciatum* | Bivalvia | Hard |
| *Castalia punctata* | Polychaeta | Easy |
| *Clupea harengus* | Actinopterygii | Hard |
| *Corophium spp.* | Malacostraca | Easy |
| *Crangon crangon* | Malacostraca | Hard |
| *Cumacea spp.* | Malacostraca | Easy |
| *Mysidacea spp.* | Malacostraca | Hard |
| *Cyprina islandica* | Bivalvia | Hard |
| *Diastylis rathkei* | Malacostraca | Easy |
| *Disoma multisectosum* | Polychaeta | Easy |
| *Euchone papillosa* | Polychaeta | Easy |
| *Gastrosaccus spinifer* | Malacostraca | Hard |
| *Gobiidae spp.* | Actinopterygii | Hard |
| *Halicryptus spinolosus* | Halicryptomorpha | Hard |
| *Harmothoe imbricata* | Polychaeta | Easy |
| *Harmothoe spp.* | Polychaeta | Easy |
| *Hyperia galba* | Malacostraca | Easy |
| *Idothea spp.* | Malacostraca | Hard |
| *Isopoda spp.* | Malacostraca | Hard |
| *Limanda limanda* | Actinopterygii | Hard |

| *Prey species* | Class | Digestibility |
|---|---|---|
| *Macoma spp.* | Bivalvia | Hard |
| *Metridium senile* | Anthozoa | Hard |
| *Microdeutopus sp.* | Malacostraca | Easy |
| *Musculus spp.* | Bivalvia | Hard |
| *Mya truncata, Mya arenaria* | Bivalvia | Hard |
| *Mysis mixta* | Malacostraca | Hard |
| *Mytilus edulis* | Bivalvia | Hard |
| *Nemertea spp.* | Nemertea | Easy |
| *Nephthys spp.* | Polychaeta | Easy |
| *Nucula nitida* | Bivalvia | Hard |
| *Ophiura albida* | Ophiuroidea | Hard |
| *Other Decapoda* | Decapoda | Hard |
| *Other Gastropoda* | Gastropoda | Hard |
| *Other Polychaeta* | Polychaeta | Easy |
| *Pectinaria koreni* | Polychaeta | Easy |
| *Phaxas pellucidus* | Bivalvia | Hard |
| *Pherusa plumosa* | Polychaeta | Easy |
| *Phtisica marina, Caprella* | Malacostraca | Easy |
| *Pisces spp.* | Actinopterygii | Hard |
| *Pleuronectiformes spp.* | Actinopterygii | Hard |
| *Polydora sp.* | Polychaeta | Easy |
| *Pomatoschistus minutus* | Actinopterygii | Hard |
| *Priapulus caudatus* | Priapulida | Easy |
| *Saxicava arctica* | Bivalvia | Hard |
| *Scoloplos armiger* | Polychaeta | Easy |
| *Spionidae spp.* | Polychaeta | Easy |
| *Terebellides stroemi* | Polychaeta | Easy |
| *Thyonidium pellucidum* | Holothuroidea | Hard |

### Supplementary information IX. Definition of default model parameters

**Table SI IX.1**: values and units of variables as set by default with the ATNr the package for the unscaled with nutrient version

| Variable (units) | parameter used | values |
|---|---|---|
| $b_{ij}$ $(m^2.s^{-1})$ | $b_0$ | 50 |
| | $b_1$ | $\mathcal{N}(0.15{,}0.03)$ |
| | $b_2$ | $\mathcal{N}(0.47{,}0.04)$ |
| | $E_a$ | -0.38 |
| $h_{ij}$ $(s)$ | $h_0$ | 0.4 |
| | $h_1$ | $\mathcal{N}(-0.66{,}0.02)$ |
| | $h_2$ | $\mathcal{N}(-0.48{,}0.03)$ |
| | $E_h$ | 0.26 |
| $c_i$ $(s)$ | $c$ | $\mathcal{N}(0.8{,}0.2)$ |
| $K_{np}$ $(g.m^{-2})$ | $k$ | $\mathcal{U}(0.1{,}0.2)$ |
| $v_{np}$ $(unitless)$ | $v$ | $\mathcal{U}(1{,}2), \mid \sum_n (v_{np} = 1)$ |
| $S_n$ $(g.m^{-2})$ | $s$ | $\mathcal{N}(10{,}2)$ |
| $r_p$ $(s^{-1})$ | $r_1$ | -0.25 |
| | $E_r$ | -0.84 |
| $X_i$ $(J.s^{-1})$ | $x_0$ | 0.138 ($i$ is a plant) |
| | $x_0$ | 0.314 ($i$ is an animal) |
| | $x_1$ | -0.25 |
| | $E_x$ | -0.69 |
| $q_i$ | - | 0.2 |
| $D$ | - | 0.25 |
